# Recurrent interactions in local cortical circuits

**DOI:** 10.1101/822700

**Authors:** Simon Peron, Ravi Pancholi, Bettina Voelcker, Jason D. Wittenbach, H. Freyja Ólafsdóttir, Jeremy Freeman, Karel Svoboda

## Abstract

The majority of cortical synapses are local and excitatory. Local recurrent circuits could implement amplification, allowing for pattern completion and other computations^1^. Cortical circuits contain subnetworks, consisting of neurons with similar receptive fields and elevated connectivity relative to the network average^2,3^. Understanding the computations performed by these subnetworks during behavior has been hampered by the fact that cortical neurons encoding different types of information are spatially intermingled and distributed over large brain volumes ^4,5^. We used computational modeling, optical recordings and manipulations to probe the function of recurrent coupling in layer (L) 2/3 of the somatosensory cortex during tactile discrimination. A model of L2/3 dynamics revealed that recurrent excitation enhances sensory signals via amplification, but only for subnetwork with elevated connectivity. Networks with high amplification were sensitive to damage: loss of a few subnetwork members degraded stimulus encoding. We tested this prediction experimentally by mapping neuronal selectivity^5^ and photoablating^6,7^ neurons with specific selectivity. In L2/3 of the somatosensory cortex, ablating a small proportion (10-20, < 5 % of the total) of neurons representing touch dramatically reduced responses in the spared touch representation, but not other representations. Network models further predicted that degradation following ablation should be greatest among spared neurons with stimulus responses that were most similar to the ablated population. Consistent with this prediction, ablations most strongly impacted neurons with selectivity similar to the ablated population. Our data shows that recurrence among cortical neurons with similar selectivity can drive input-specific amplification during behavior.

---

Two circuit motifs have been proposed to account for dynamics in cortical layer (L) 2/3: recurrent excitation, which may cause amplification^1,8^, and feedback inhibition, which may account for the sparse activity typically observed in L2/3^9,10^. We explored the role of these motifs in a model of layer (L) 2/3 of the vibrissal somatosensory cortex (vS1). The model is constrained by measured neuronal biophysical parameters, connection probabilities, and connection strengths^11,12^ (Methods). The network included 1,700 excitatory and 300 inhibitory neurons, implemented as integrate-and-fire cortical neurons.

To model input-specific recurrent coupling, we restricted the sensory input to a subnetwork of the excitatory neurons (200 of 1,700), corresponding to the number of neurons representing touch in L2/3 of vS1 during active tactile behavior^5,13^ (Fig. 1a). L2/3 neurons receive input from L4^12^. We simulated touch-related input to L2/3 based on L4 recordings^14^ (latency of peak response, 10 ms; full width half-max, FWHM, 12.8 ms).

**Figure 1.**
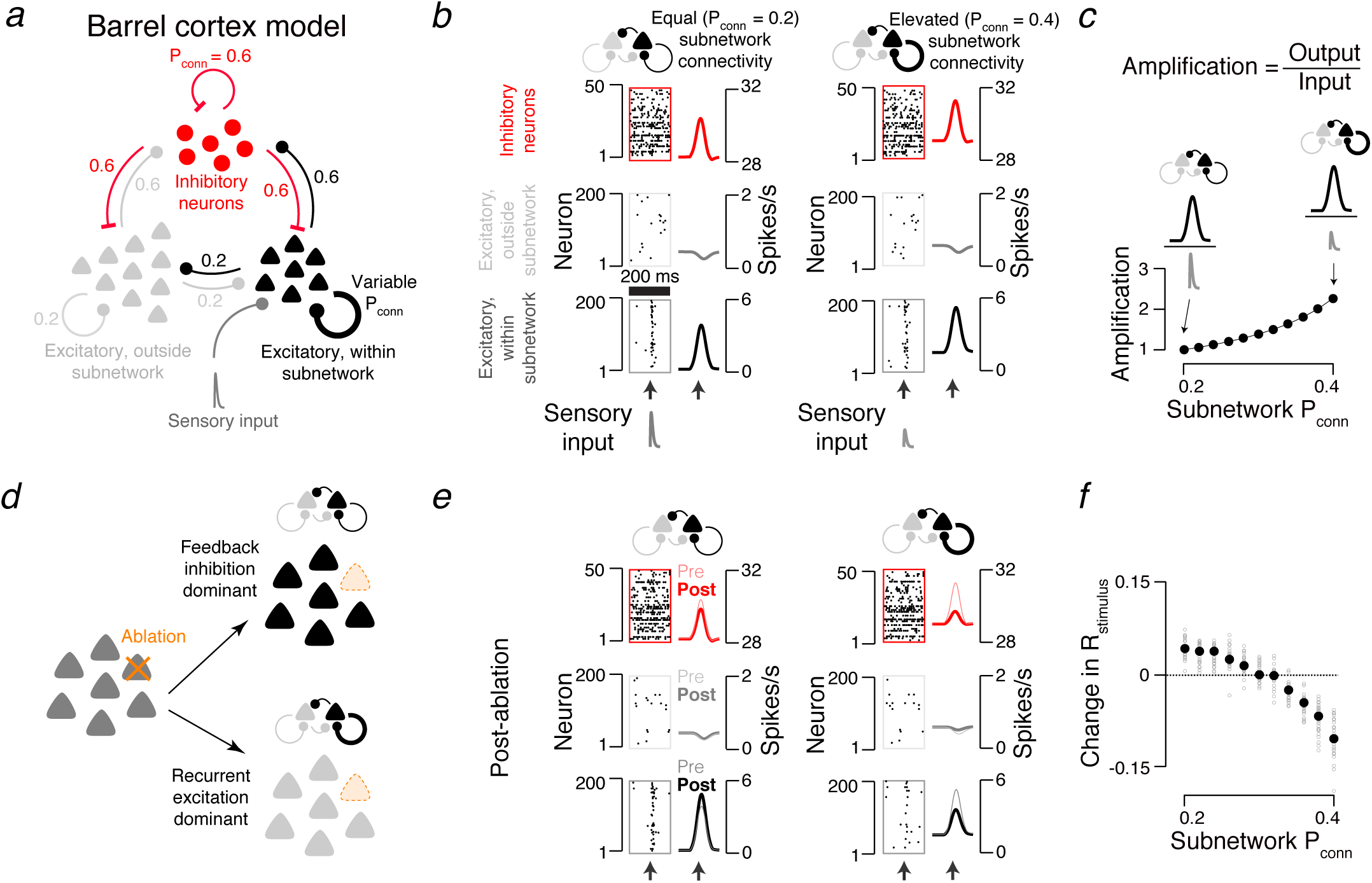
Ablation effect in simulated cortical L2/3 network depends on excitatory connectivity. **a.** The network model of layer 2/3 comprises a small subnetwork of 200 excitatory neurons (black) receiving direct sensory input (dark grey), a larger excitatory population (1,500 neurons; light grey) without direct input, and 300 inhibitory neurons (red). All populations are interconnected (connection probabilities, P_conn_, listed in figure; Methods). The probability and strength of connections within the excitatory subnetwork was varied (thick black loop; Methods). **b.** Model network responses aligned to input (arrow). Left, subnetwork connectivity for excitatory subnetwork equal to overall connection probability. Right, elevated subnetwork connectivity. Raster plots show a subset of neurons from an example network. Peri-stimulus time histograms show averages across all neurons and networks. Bottom, excitatory neurons within subnetwork; middle, excitatory neurons outside subnetwork; top, inhibitory neurons. **c.** Amplification, defined as the ratio of network output to sensory input (Methods), as a function of subnetwork connectivity, normalized to the P_conn_ = 0.2 case.. **d.** Prediction of the effect of ablating a small number of neurons in the two architectures. In the equal connectivity case (top), feedback inhibition dominates and responses among spared neurons increase. In the elevated connectivity case (bottom), recurrent excitation dominates and responses decline. **e**. As in **b**, but following ablation. Pre-ablation peri-stimulus time histograms are shown as thinner lines. **f.** Effect of ablating the 20 neurons with the strongest encoding score on stimulus encoding, as a function of subnetwork connectivity. Grey dots, individual simulations. Black circles, mean of 30 simulated networks.

To measure the faithfulness with which the spike rate of a neuron reflects the sensory input, we computed an ‘encoding score’ (R_stimulus_) by cross-correlating the activity of each neuron with the input (Methods).

We varied recurrence by changing the connection probability within the input-recipient subnetwork (‘subnetwork connectivity’; synaptic conductance was scaled proportionately 15, Methods). For each subnetwork connectivity we matched the encoding score distribution to that observed *in vivo*^5^ by adjusting the strength of the input from L4 (Methods). Subnetworks with connectivity equal to and moderately elevated relative to the rest of the network (non-subnetwork connectivity, 0.2; subnetwork connectivity range, 0.2 - 0.4) produced responses consistent with those observed *in vivo*^5^ (Fig. 1b; Methods). The amplitudes of the required sensory input declined with increasing subnetwork connectivity. Amplification, defined as the ratio of network output to network input, therefore increased with subnetwork connectivity^16^ (Fig. 1c). Additional elevation of subnetwork connectivity (> 0.4) produced all-or-none network responses, whereby a transient input drove the network into a persistently active state, as in models of memory-related activity in frontal cortex ^17^ (Extended Data 1).

Overall, subnetwork behavior fell into three regimes, each of which produced a distinct response to removal (‘ablation’) of a small number of constituent elements. Subnetworks with low connectivity (0.2) amplified little, and were resistant to ablation of parts of the network (Fig. 1c-f). Encoding scores for spared neurons increased following simulated ablation of 10 % of the subnetwork, because of reduced competition by feedback inhibition^9,18^ (encoding score, from 0.237 ± 0.027 to 0.274 ± 0.032, grand median encoding score ± adjusted median absolute deviation, MAD, P < 0.001, Wilcoxon signed rank test, across N = 30 simulated networks with different randomized connectivity and initial conditions; Methods) (Fig. 1d-f).

Subnetworks with elevated connectivity (e.g., 0.4) exhibited stronger amplification^16,19^ (Fig. 1c). Ablations now caused encoding scores to decline for spared neurons, from 0.243 ± 0.049 to 0.143 ± 0.028 (connectivity = 0.4; P < 0.001, N = 30 networks), because of reduced nonlinear amplification (Fig. 1d-f). Networks in the all-or-none regime^17^ were robust to ablation, maintaining their all-or-none response (Extended Data 1). The response of subnetworks to ablations therefore distinguishes the three network regimes.

We next performed a similar analysis for actual L2/3 networks during behavior. Mice were trained to localize an object with one spared whisker (Fig. 2a)^5^. Behavioral trials contained a sample epoch (duration, 1 s), during which mice touched a vertical pole presented in one of a range of locations (Fig. 2a, b), a delay epoch (0.5-1 s) during which mice withheld their response, followed by a response epoch, during which mice reported pole position by licking a left or right ‘lickport’.

**Figure 2.**
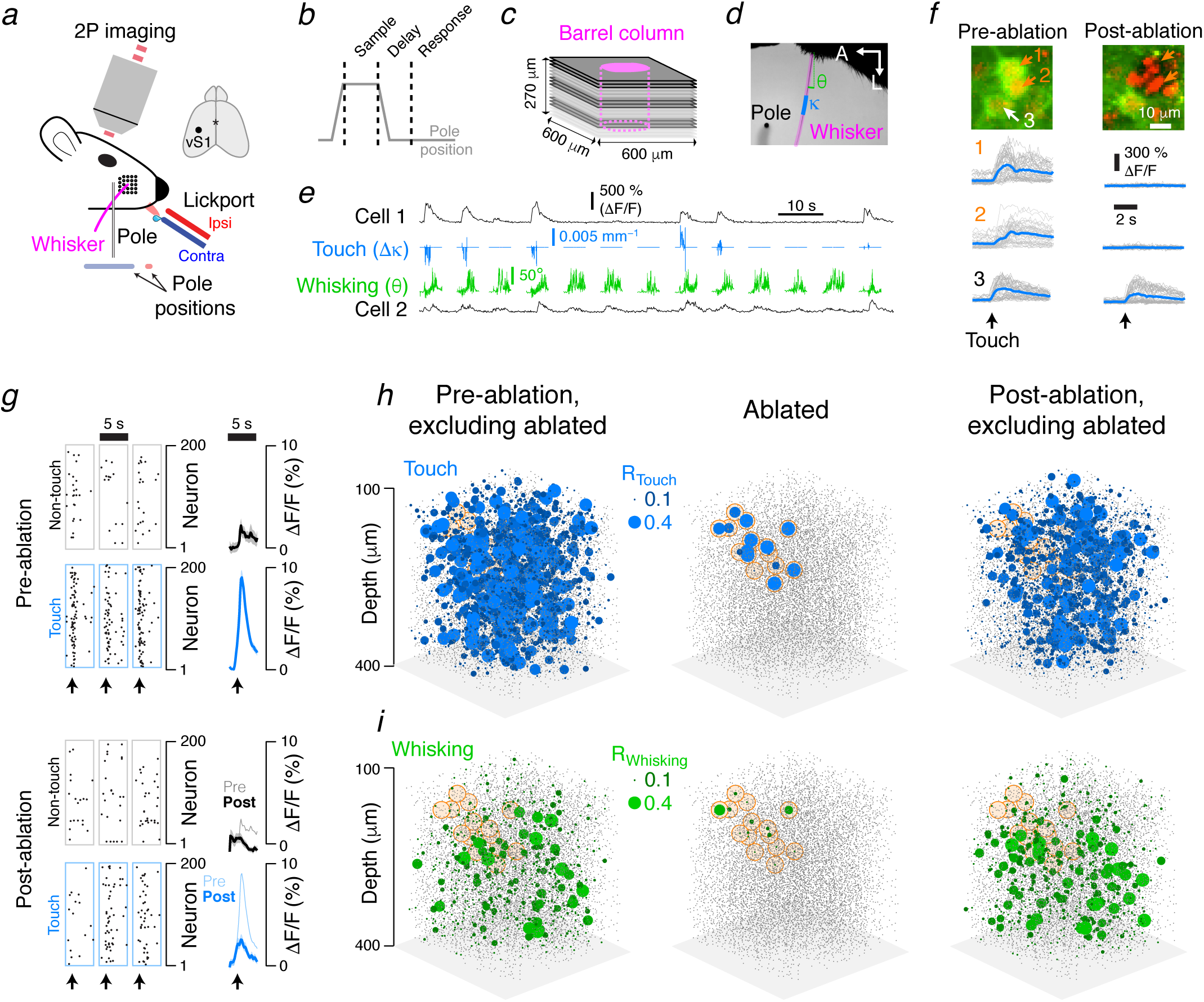
Vibrissal S1 touch networks are sensitive to ablation. **a.** Mice used one whisker to discriminate the position of a pole and reported perceived position by licking the left (ipsilateral with respect to side of brain imaged, red) or right (contralateral, blue) lickport. **b.** Task epochs. The pole (grey line, **a**) was accessible during the sample epoch. An auditory cue initiated the response epoch. **c.** Imaging. Imaging planes (600 x 600 µm) were centered on the barrel column of the spared whisker, magenta. Three planes (15 µm apart) were acquired simultaneously for ∼5-10 minutes (20-60 trials), after which the next set of three (same shade of grey) was imaged. **d.** Example video frame showing the whisker (magenta), pole (black, bottom-left), whisker curvature (κ, blue), and whisker angle (θ, green). **e.** Example neuronal activity for a touch (cell 1, top trace) and whisking (cell 2, bottom trace) cell. Blue, whisker curvature change (Δκ, ‘touch’); green, whisker angle (θ, ‘whisking’). **f.** Top, three pyramidal neurons prior to (left) and the day after (right) ablation. Green, GCaMP6s; red, mCherry. Neurons 1 and 2 (orange) were ablated; neuron 3 was spared. Bottom, touch-aligned neuronal responses before (left) and after (right) ablation. **g-i.** Example experiment. **g.** Example touch responses (dots, calcium events; Methods) for a subset of neurons before (top) and after (bottom) ablation. Grey boxes, 200 non-touch/non-whisking neurons; blue boxes, 200 touch neurons. Example touches were selected so that the forces of pre- and post-ablation touches were similar. Right, touch-aligned mean ΔF/F averaged across these same touch (blue) and non-touch/non-whisking (grey) neurons across the strongest 5% of touches before (top) and after (bottom) ablation. Envelopes show standard error of the mean. **h.** Left, example map for touch cells before ablation. Sphere size corresponds to R_touch_ (top). Grey dots, other neurons. The map excludes the ablated neurons, whose position is indicated with a faint orange background. Center, R_touch_ for the ablated population. Right, R_touch_ following ablation, with ablated neurons again excluded. **i.** As for **h**, but for whisking neurons.

We imaged activity in vS1^5^. L2/3 neurons (8,164 ± 2,569 per mouse, mean ± S.D.; N = 16 mice; Extended Data 2) were sampled in the barrel column corresponding to the spared whisker using volumetric imaging (Methods; Fig. 2c). Activity in a subset of neurons encoded whisker position (‘whisking’ neurons), whereas others responded to touch-induced changes in whisker curvature (‘touch’ neurons)^5^ (Fig. 2d, e). We quantified the relationship of specific behavioral features and neural activity. An encoding model generated a prediction of neural activity. The correlation of the prediction with the actual neural activity comprised an ‘encoding score’ (Methods), which was used to classify neurons as belonging to the touch and/or the whisking representation^5^ (Methods). Neurons representing touch and whisking were spatially intermingled. Across mice (N = 16 mice; Extended Data 1), 961 ± 660 neurons of the imaged neurons encoded touch (range: 152 to 2,338; fraction: 0.113 +/-0.062) and 892 ± 408 encoded whisker movements (range: 348 to 1,516; fraction: 0.108 +/-0.030).

We probed the role of recurrence by ablating members of the touch representation and examining the effect on spared neurons. Groups of individual excitatory neurons were ablated using multiphoton excitation^6,7,20,21^ (Extended Data 3, 4).

Ablating a small proportion of strong touch cells (Fig. 2f; 16.8 ± 12.8 neurons, 6 % of touch neurons in the spared whisker’s barrel column, N = 9 mice; touch score, 70^th^ ± 35^th^ percentile; Extended Data 1) reduced responses to touch in the spared touch neurons (Fig. 2g, h). The touch encoding score (R_touch_) across touch neurons declined (from 0.123 ± 0.021 to 0.100 ± 0.037, grand median ± adjusted MAD; N = 9 mice, 8,392 neurons, P = 0.004, Wilcoxon signed rank test; Fig. 3a; Methods), as did the touch neuron count (from 932 ± 634 to 716 ± 469, mean ± SD, calculations exclude ablated neurons, Methods). The whisking encoding score (R_whisking_) did not change (Fig. 2i, 3b; from 0.116 ± 0.013 to 0.115 ± 0.024; 6,975 neurons, P = 0.820; neuron count: from 775 ± 267 to 721 ± 241).

**Figure 3.**
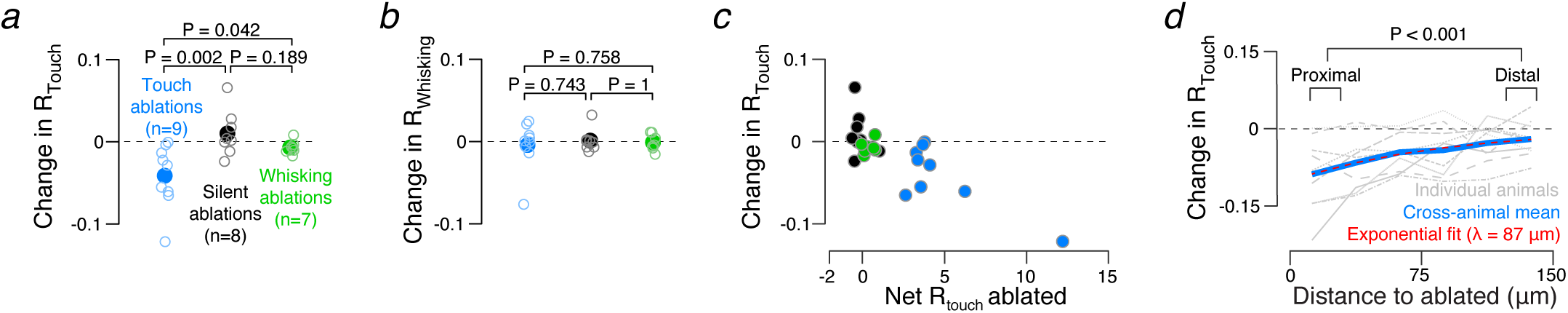
Parameters influencing ablation sensitivity. **a.** Effect of ablating different populations on R_touch_. Blue, touch ablations; black, silent neuron ablations; green, whisking neuron ablations. Only touch ablations produced a significant effect on R_touch_ (see text). **b.** Effect of ablating different populations on R_whisking_; as in **a. c.** Ablation effect on touch representation (quantified as change in R_touch_) depends on the net amount of touch representation degradation (total R_touch_ across ablated neurons). Color scheme as in **a, b. d.** Effect of touch neuron ablation on R_touch_ as a function of distance from an ablated neuron; grey, individual mice; blue, cross-animal mean; stippled red, exponential fit.

Our model predicts that the decline following ablation should be proportional to the extent of the network removed (Extended Data 5). Consistent with this prediction, the decline in touch representation increased as more of the touch representation was ablated (Fig. 3c; Pearson correlation of change in R_touch_ and net R_touch_ ablated, R = −0.80, P < 0.001 across all 24 ablations; R = −0.78, P = 0.013 when restricted to the 9 touch cell ablations). Touch neurons more proximal (15-35 µm) to the ablated cells experience a larger decline in R_touch_ than those distal (115-135 µm) from the ablated cells (proximal change in ΔR_touch_: −0.132 ± 0.246, grand median ± adjusted MAD, distal: −0.056 ± 0.241, N = 9 mice, P = 0.020, Wilcoxon signed rank test comparing proximal to distal, paired by animal). Thus, the effects of ablating the touch neurons decayed over a distance (λ = 87 µm, exponential fit; Fig. 3d) that is similar to the spatial scale of local recurrent connectivity in the rodent sensory cortex^22^. No distance-dependence was observed following touch ablation among whisking neurons (Extended Data 6). The declining touch representation was not caused by fewer or weaker touches, or different patterns of whisker movements (Extended Data 7). This result is consistent with amplification of touch responses by recurrent excitation in L2/3.

In contrast, ablating a subset of strong whisking neurons (12.7 ± 5.7 neurons, ∼4 % of whisking neurons in the spared whisker’s barrel column, N = 7 mice; whisking score: 68^th^ ± 26^th^ percentile) produced no effect on either the touch (Extended Data 8, Fig. 3a; R_touch_ from 0.110 ± 0.080 to 0.095 ± 0.069, N = 7 mice, 5,899 neurons, P = 0.109, count from 843 ± 403 to 878 ± 540) or the whisking representation (R_whisking_ from 0.108 ± 0.076 to 0.107 ± 0.076, N = 6,395, P = 0.812, count from 914 ± 434 to 970 ± 554). Similarly, ablating silent neurons (event rate below 0.025 Hz; Methods; 16.3 ± 2.6 neurons, N = 8 mice) did not change the touch (Extended Data 8, Fig. 3b; R_touch_ from 0.115 ± 0.021 to 0.107 ± 0.024, N = 8 mice, 7,110 neurons, P = 0.383, count from 889 ± 713 to 926 ± 681) or whisking representations (R_whisking_ from 0.115 ± 0.014 to 0.114 ± 0.015, 7,684 neurons, P = 0.844, count from 961 ± 464 to 904 ± 384). Touch ablations produced significantly different encoding score changes among touch (but not whisking) representations than either whisking or silent ablations (Fig. 3a,b; touch vs. whisking ablations, touch network, P = 0.042; whisking network, P = 0.758; touch vs. silent ablations, touch network, P = 0.002; whisking network, P = 0.743; Wilcoxon rank sum test, Methods), whereas whisking ablations did not produce significantly different changes from silent ablations (touch network, P = 0.189; whisking network, P = 1.000; Wilcoxon rank sum test, Methods). Non-specific effects of ablation therefore do not contribute to changes in the touch representation after ablation.

Ablations of whisking neurons did not degrade the representation of whisking. Our network model suggests that the slower kinetics of the whisking input alone does not account for the lack of ablation effect, as networks having elevated subnetwork connectivity (0.4) with slower input (t_peak_ = 50 ms, vs. 10 ms for touch simulations; Methods) were sensitive to simulated ablation (encoding score change after ablation: from 0.143 ± 0.024 to 0.104 ± 0.013, P < 0.001, N = 30 networks; Extended Data 9). However, individual neurons encode whisking input with different phases^23^, which suggests an asynchronous population response that would be unlikely to engage recurrent excitation. In contrast, rapid touch input is in-phase across the neural population^13,14^. Therefore, the lack of sensitivity to ablation in the whisking population does not necessarily imply an absence of effective recurrent coupling.

The responses of individual neurons differ in complex ways not captured by the binary classification scheme (‘touch’ or ‘whisking’). For example, touch neurons show directional preference and diverse response dynamics^5^. In recurrent networks connected in a feature-specific manner^15^, the effects of ablation on spared neurons should increase with the similarity of their tuning to the ablated population^9,24,25^.

We tested this intuition in our L2/3 model, defining ‘response similarity’ as the correlation of a neuron’s activity with the mean activity of the ablated neurons (Fig. 4a; Methods). In networks with elevated subnetwork connectivity, neurons with high response similarity showed the largest decline in encoding score following ablation (Fig. 4b; P = 0.200, sign test, testing that the slope of change in encoding score as a function of response similarity is 0, N = 30 networks, Methods; 3,466 neurons shown). In networks without elevated subnetwork connectivity, the relationship disappeared (P = 0.001, N = 30 networks; 3,779 neurons shown).

**Figure 4.**
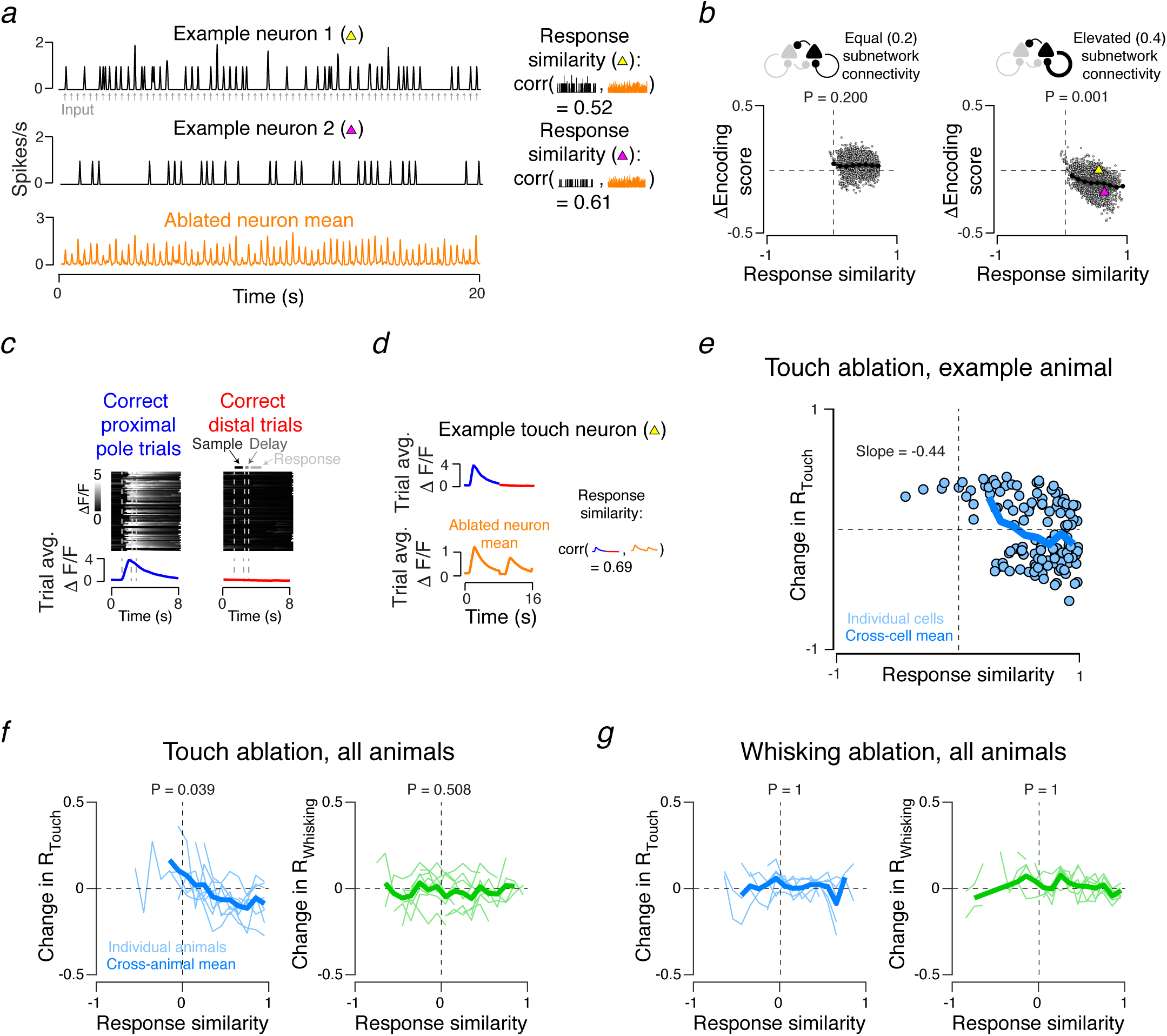
Effect of ablations depends on response similarity to ablated cells. **a.** Computing response similarity for individual neurons in simulated networks. Spike rate for each neuron (black, top two traces) was obtained by convolving spiking response with a Gaussian (σ = 20 ms; Methods). Grey arrows, sensory input. Orange, mean spike rate across ablated neurons. Right, response similarity was computed for individual neurons by correlating spike rate with the mean spike rate of the ablated neurons. **b.** In model networks with elevated subnetwork connectivity the post-ablation encoding score change depends on response similarity. Left, network where subnetwork connectivity equaled remaining excitatory connectivity. Right, network with elevated subnetwork connectivity. Grey dots, individual neurons. Dark grey lines, single network averages; p-values are given for the sign test evaluating whether the slopes of individual network linear fits (N=30) are 0. Black line, grand mean across networks. Yellow and magenta triangles (right), neurons shown in **a. c.** Computing the trial averaged response for a vS1 touch neuron. Heat maps show ΔF/F for individual trials (correct contralateral lick trials, left; correct ipsilateral, right). Sample, delay, and response epochs (Fig. 1b) are delineated by dashed lines. Bottom, trial averaged ΔF/F for both trial types. **d.** Top, trial averaged ΔF/F for neuron in **c**. Individual neuron trial averaged response vectors are constructed by concatenating the averaged response for the two trial types. Bottom, the mean trial averaged response across ablated neurons. Right, correlating the individual neuron response vector with the mean across ablated neurons gives the response similarity. **e.** Touch neuron ablation induced changes in encoding model touch score as a function of response similarity in an example animal. Blue circles, individual neurons; blue line, mean for this animal. Slope for a linear fit of change in R_touch_ as a function of response similarity is given. **f.** Population data for all touch ablations for both touch (blue, left) and whisking (right, green) neurons. Thin colored lines, individual animal means for a given response similarity bin; thick lines, cross-animal mean. P-values for sign test that the slope of change in encoding score as a function of response similarity is 0 are given (N=9). **g**. As in **f**, but for whisking neuron ablation. P-values for sign test as in **f** (N=7).

We next performed a similar analysis on our experimental data. Our imaging experiments did not track all neurons simultaneously (we imaged 3/18 planes at a time; Fig. 2c; Methods) ^5^. We therefore devised an alternative ‘response similarity’ metric that did not depend on simultaneous recording. For each neuron, we averaged responses across trials for both trial types (correct contra and ipsi trials; Fig. 4c) prior to ablation. Concatenating the contra and ipsi responses yielded a trial-averaged ΔF/F for each neuron (Fig. 4d). Computing the mean trial-averaged ΔF/F across all ablated neurons provided the ablated neuron mean response. Response similarity was measured for each neuron as the correlation between its trial-averaged ΔF/F response and the mean response across all ablated neurons.

Changes in R_touch_ following ablation depended on response similarity (Fig. 4e, f; P = 0.039, sign test, testing that the slope of change in encoding score as a function of response similarity is 0, N = 9 mice; 2,770 neurons total; Methods). R_touch_ in neurons with high response similarity declined. Neurons with negative response similarity showed increased R_touch_, potentially due to reduced feedback inhibition previously evoked by the ablated neurons ^9^ (Fig. 4e).

Touch neuron ablation had no effect on the whisking network (Fig. 4f; P = 0.508, sign test, N = 9 mice; 1,105 neurons). Whisker neuron ablation had no effect on either representation (Fig. 4g; N = 7 mice, touch: P = 1; 1,639 neurons total; whisking: P = 1; 959 neurons total).

Targeted photoablation in cortical networks allowed us to test roles of effective recurrence in cortical circuits ^1^. The selective degradation of representations resembling the ablated neurons is consistent with strong amplification in recurrent networks^16,24,26^ (Fig. 1), but inconsistent with networks dominated by feedback inhibition^9,19,27^ or stable attractor states^17,28^. Our work suggests that within a given cortical area, distinct circuitry underpin the processing of different stimulus classes. Our experiments reveal that cortical circuits can be surprisingly sensitive to damage targeted of a specific representation, despite remarkable robustness to other kinds of perturbations^29^.

## Author contributions

SP and KS conceived the project. SP, RP, and TV performed the experiments, with assistance from HO. JW performed the modeling, with input from JF, SP and KS. SP, RP, TV, JW, JF, and KS analyzed data and wrote the paper.

## Acknowledgments

We thank Shaul Druckmann, Sandro Romani, Diego Gutnisky, Nuo Li, Jianing Yu, Hidehiko Inagaki, Nicholas Sofroniew, and Mike Economo for comments on the manuscript, and Amy Hu for histology. Funding was provided by the Howard Hughes Medical Institute. RP was supported by NIH T32GM007308.

## References

1 Douglas, R. J. & Martin, K. A. Recurrent neuronal circuits in the neocortex. Curr Biol 17, R496–500, doi:10.1016/j.cub.2007.04.024 (2007).

2 Ko, H. et al. Functional specificity of local synaptic connections in neocortical networks. Nature 473, 87–91, doi:10.1038/nature09880 (2011).

3 Biagi, R., Cossellu, G., Sarcina, M., Pizzamiglio, I. T. & Farronato, G. Laser-assisted treatment of dentinal hypersensitivity: a literature review. Annali di stomatologia 6, 75–80, doi:10.11138/ads/2015.6.3.075 (2015).

4 Ohki, K. & Reid, R. C. Specificity and randomness in the visual cortex. Curr Opin Neurobiol 17, 401–407, doi:S0959-4388(07)00090-6 [pii] 10.1016/j.conb.2007.07.007 (2007).

5 Peron, S. P., Freeman, J., Iyer, V., Guo, C. & Svoboda, K. A Cellular Resolution Map of Barrel Cortex Activity during Tactile Behavior. Neuron 86, 783–799, doi:10.1016/j.neuron.2015.03.027 (2015).

6 Konig, K., Becker, T. W., Fisher, P., Riemann, I. & Halbhuber, K. J. Pulse-length dependence of cellular response to intense near-infrared laser pulses in multiphoton microscopes. Optics Lett. 24, 113–115 (1999).

7 Vladimirov, N. et al. Brain-wide circuit interrogation at the cellular level guided by online analysis of neuronal function. Nat Methods 15, 1117–1125, doi:10.1038/s41592-018-0221-x (2018).

8 London, M., Roth, A., Beeren, L., Hausser, M. & Latham, P. E. Sensitivity to perturbations in vivo implies high noise and suggests rate coding in cortex. Nature 466, 123–127, doi:10.1038/nature09086 (2010).

9 Chettih, S. N. & Harvey, C. D. Single-neuron perturbations reveal feature-specific competition in V1. Nature 567, 334–340, doi:10.1038/s41586-019-0997-6 (2019).

10 O’Connor, D. H., Peron, S. P., Huber, D. & Svoboda, K. Neural activity in barrel cortex underlying vibrissa-based object localization in mice. Neuron 67, 1048–1061, doi:10.1016/j.neuron.2010.08.026 (2010).

11 Avermann, M., Tomm, C., Mateo, C., Gerstner, W. & Petersen, C. C. Microcircuits of excitatory and inhibitory neurons in layer 2/3 of mouse barrel cortex. J Neurophysiol 107, 3116–3134, doi:10.1152/jn.00917.2011 (2012).

12 Lefort, S., Tomm, C., Floyd Sarria, J. C. & Petersen, C. C. The excitatory neuronal network of the C2 barrel column in mouse primary somatosensory cortex. Neuron 61, 301–316, doi:10.1016/j.neuron.2008.12.020 (2009).

13 Crochet, S., Poulet, J. F., Kremer, Y. & Petersen, C. C. Synaptic mechanisms underlying sparse coding of active touch. Neuron 69, 1160–1175, doi:10.1016/j.neuron.2011.02.022 (2011).

14 Hires, S. A., Gutnisky, D. A., Yu, J., O’Connor, D. H. & Svoboda, K. Low-noise encoding of active touch by layer 4 in the somatosensory cortex. Elife 4, doi:10.7554/eLife.06619 (2015).

15 Okun, M. et al. Diverse coupling of neurons to populations in sensory cortex. Nature 521, 511–515, doi:10.1038/nature14273 (2015).

16 Douglas, R. J., Koch, C., Mahowald, M., Martin, K. A. & Suarez, H. H. Recurrent excitation in neocortical circuits. Science 269, 981–985 (1995).

17 Litwin-Kumar, A. & Doiron, B. Slow dynamics and high variability in balanced cortical networks with clustered connections. Nat Neurosci 15, 1498–1505, doi:10.1038/nn.3220 (2012).

18 Barrett, D. G., Deneve, S. & Machens, C. K. Optimal compensation for neuron loss. Elife 5, doi:10.7554/eLife.12454 (2016).

19 Hansel, D. & van Vreeswijk, C. The mechanism of orientation selectivity in primary visual cortex without a functional map. J Neurosci 32, 4049–4064, doi:10.1523/JNEUROSCI.6284-11.2012 (2012).

20 Orger, M. B., Kampff, A. R., Severi, K. E., Bollmann, J. H. & Engert, F. Control of visually guided behavior by distinct populations of spinal projection neurons. Nat Neurosci 11, 327–333, doi:10.1038/nn2048 (2008).

21 Allegra Mascaro, A. L., Sacconi, L. & Pavone, F. S. Multi-photon nanosurgery in live brain. Frontiers in neuroenergetics 2, doi:10.3389/fnene.2010.00021 (2010).

22 Holmgren, C., Harkany, T., Svennenfors, B. & Zilberter, Y. Pyramidal cell communication within local networks in layer 2/3 of rat neocortex. J Physiol 551, 139–153, doi:10.1113/jphysiol.2003.044784 (2003).

23 Curtis, J. C. & Kleinfeld, D. Phase-to-rate transformations encode touch in cortical neurons of a scanning sensorimotor system. Nat Neurosci 12, 492–501 (2009).

24 Marshel, J. H. et al. Cortical layer-specific critical dynamics triggering perception. Science 365, doi:10.1126/science.aaw5202 (2019).

25 Carrillo-Reid, L., Han, S., Yang, W., Akrouh, A. & Yuste, R. Controlling Visually Guided Behavior by Holographic Recalling of Cortical Ensembles. Cell 178, 447–457 e445, doi:10.1016/j.cell.2019.05.045 (2019).

26 Lien, A. D. & Scanziani, M. Tuned thalamic excitation is amplified by visual cortical circuits. Nat Neurosci 16, 1315–1323, doi:10.1038/nn.3488 (2013).

27 Barth, A. L. & Poulet, J. F. Experimental evidence for sparse firing in the neocortex. Trends Neurosci 35, 345–355, doi:10.1016/j.tins.2012.03.008 (2012).

28 Wang, X. J. Decision making in recurrent neuronal circuits. Neuron 60, 215–234 (2008).

29 Li, N., Daie, K., Svoboda, K. & Druckmann, S. Robust neuronal dynamics in premotor cortex during motor planning. Nature, doi:10.1038/nature17643 (2016).

