## Supplementary Information for "Recurrent interactions in local cortical circuits"

#### Supplementary Materials

| Animal ID | Cell count | Ablated type | Ablated count | Touch count | $\Delta R_{touch}$ | Whisking count | $\Delta R_{whisking}$ |
|---|---|---|---|---|---|---|---|
| **j250220** | 8,844 | vS1 touch | 53 | 867 | -0.122 | 422 | -0.076 |
| **j257218** | 6,215 | vS1 silent | 20 | 743 | 0.018 | 366 | 0.032 |
| **j257220** | 3,663 | vS1 touch | 9 | 389 | -0.004 | 455 | 0.025 |
| **j258836** | 7,290 | vS1 silent | 16 | 481 | 0.066 | 666 | -0.012 |
| | | vS1 touch | 15 | 488 | -0.055 | 655 | 0.001 |
| **j271211** | 9,415 | vS1 silent | 16 | 2,476 | -0.012 | 551 | -0.007 |
| | | vS1 touch | 15 | 1,951 | -0.061 | 692 | 0.021 |
| **j278937** | 4,230 | vS1 silent | 15 | 146 | 0.004 | 379 | 0.001 |
| | | vS1 whisking | 12 | 150 | -0.006 | 392 | 0.011 |
| **j278939** | 7,167 | vS1 silent | 20 | 636 | -0.024 | 698 | 0.001 |
| | | vS1 whisking | 13 | 523 | -0.017 | 649 | -0.015 |
| **j281915** | 7,002 | vS1 whisking | 8 | 723 | -0.010 | 550 | 0.004 |
| | | vS1 touch | 11 | 601 | -0.022 | 672 | 0.008 |
| **n274424** | 7,785 | vS1 touch | 17 | 706 | -0.029 | 718 | -0.014 |
| **n272761** | 6,359 | vS1 touch | 14 | 509 | -0.001 | 410 | 0.000 |
| **n274577** | 12,173 | vS1 touch | 11 | 1,380 | -0.013 | 657 | 0.004 |
| **n275801** | 8,354 | vS1 silent | 14 | 220 | 0.000 | 738 | -0.007 |
| | | vS1 touch | 9 | 197 | -0.065 | 698 | -0.006 |
| **n275798** | 11,364 | vS1 silent | 13 | 972 | 0.008 | 958 | 0.002 |
| | | vS1 whisking | 11 | 959 | 0.002 | 1,037 | 0.011 |
| **n278288** | 9,314 | vS1 whisking | 25 | 969 | -0.008 | 813 | -0.006 |
| **n276013** | 11,659 | vS1 silent | 16 | 652 | 0.028 | 1,072 | -0.001 |
| | | vS1 whisking | 9 | 597 | -0.012 | 1,228 | -0.008 |
| **n278759** | 9,187 | vS1 whisking | 11 | 617 | -0.003 | 526 | -0.004 |

**Extended Data 2.** Individual animals. Each mouse is listed along with the number of neurons imaged, the type(s) of ablation(s) performed, and the number of neurons ablated. For vS1 ablations, the number of touch and whisking neurons is given, as well as the change in median encoding score following ablation. Animals whose ID starts with a ‘j’ were virally transfected Emx1-Cre X LSL-H2B-mCherry[1]; animals whose ID starts with an ‘n’ were Ai162 X Slc17a7-Cre[2].

**Supplementary Figure Legends**

**Extended Data 1.** Effect of ablation in a simulated network with hyper-connectivity (i.e., $P_{conn} > 0.4$; here, $P_{conn} = 0.44$). Model network responses aligned to input (arrow, bottom). Raster plots show a subset of neurons from an example network. Peri-stimulus time histograms show averages across all neurons and networks. Bottom, excitatory neurons within subnetwork; middle, excitatory neurons outside subnetwork; top, inhibitory neurons. Left, network response before ablation of 10% of the subnetwork neurons; right, response after ablation.

**Extended Data 3.** Multiphoton ablation protocol. **a.** Example field of view immediately before (left) and after (right) a successful ablation. Orange arrow, target neuron. **b.** Ablation protocol. Power (orange, top trace) during ablation epochs (100 ms; elevated power, orange, top) gradually increased, and the PMT shutter was closed (black bars). During the intervening evaluation epochs, power was lower and constant (orange, top), and the PMT shutter was open. Ablation was terminated when GCaMP fluorescence at the target neuron (green) jumped (orange arrow). **c.** Success of ablation as a function of neuron depth for all experiments included in this dataset. Individual points give mean success rate for given depth bin; bin size, 25 μm. **d**. Depth dependence of total energy deposition for successful ablations (successfully ablated cells only: N = 293 cells across 24 sessions, 16 animals). Grey dots, individual ablations. Black dots, means across 25 μm bins. **e**. As in **d**, but for peak power needed for ablation (N = 293 cells across 24 sessions, 16 animals).

**Extended Data 4.** Controlled ablation produces spatially localized effects. **a.** Change in calcium event rate (Methods) as a function of minimal distance to an ablated neuron following silent cell ablation. Individual neurons appear as gray points, with dark gray dashed lines showing single animal averages and the dark black line showing the cross-animal (N = 8 silent ablation mice) average. Event rates among neurons adjacent to ablated silent cells did not change (see text). **b.** Confocal *ex vivo* image from an animal perfused 24h following ablation. Ablation sites are indicated with dashed white circles. Green, GCaMP6s fluorescence; red, mCherry fluorescence; blue, microglial antibody iba1 fluorescence. **c.** As in **b**, but blue shows immunoreactivity for the astrocytic marker, GFAP. **d.** The spatial extent of glial reaction was measured by detecting the fastest intensity decline ridge (dashed white line; Methods) in the glial antibody image along lines emanating from the ablation center at varied angles. Top, ridge along ablation from **b**. Bottom left, intensity image in angle-distance space within which ridge was measured. Bottom right, distribution of reaction radii; all points constitute iba1 labeling, as no detectable glial reactions were observed with GFAP: 8 of 10 iba1-labeled and 0 of 6 GFAP-labeled ablations retrieved histologically revealed a detectable glial reaction (Methods). These reactions had radii of 11.7 ± 1.7 μm (mean ± S.D.; N = 8). **e**. Two-photon *in vivo* images before (left) and after (right) a successful ablation (target, white dotted line). Green, GCaMP6s fluorescence; red, mCherry fluorescence. **f.** As in **e**, but following a failure of the ablation protocol to terminate the ablation. Excess energy deposition produced a large lesion (black, center of image). White arrows, corresponding points in the two images. Animals (N = 5) with such lesions were excluded from the study.

**Extended Data 5.** Ablation's effect on L2/3 model sensory representation increases with the number of ablated neurons. Ablation effect (change in $R_{touch}$) as a function of the degree of touch representation degradation (net $R_{touch}$ across ablated neurons). In all cases, we used elevated subnetwork connectivity (0.4; Figure 1, Methods). The $N_{ablated}$ neurons with the highest encoding score were selected for ablation. Grey circles, individual networks. Black dots, median across N = 30 simulated networks for a given number of ablated neurons, indicated in the plot. Beyond $N_{ablated} > 50$, we observed instability, presumably because we did not attempt to restore excitatory-inhibitory balance following ablation; these data were omitted. Correlation of net $R_{touch}$ ablated and $\Delta R_{stimulus}$: R = -0.65, P < 0.01.

**Extended Data 6.** Touch neuron ablation does not produce distance-dependent effects in the whisking representation. Proximal (15-35 μm to nearest ablated cell) change in $R_{whisking}$: 0.000 ± 0.192; distal (115-135 μm): -0.023 ± 0.186, (P = 0. 098, Wilcoxon signed rank test comparing proximal paired to distal, N = 9 mice); legend as in Fig. **3d**.

**Extended Data 7.** Behavior does not account for ablation effects. **a.** Pre- (dark blue) and post-touch neuron ablation (light blue) distributions for whisker angle ( $\theta$ ), angle at touch ( $\theta$ at touch), and net curvature change across all touches (net $\Delta\kappa$ at touch) in an example animal. P-values, Kolmogorov-Smirnov test. No animal showed a significant (P < 0.05) change in any of the three parameters. **b.** Fraction of correct trials (left) and number of touches (right) before and after touch neuron ablation. P-values, Wilcoxon signed rank test, N = 9 mice. **c, d.** As in **a, b**, but for silent neuron ablations, N = 8 mice. **e, f.** As in **a, b**, but for whisking neuron ablations, N = 7 mice.

**Extended Data 8.** Example effects of whisking and silent neuron ablations. **a**. Example whisking neuron ablation. Left, example maps for touch (top, blue) and whisking (bottom, green) cells before ablation. Sphere size corresponds to $R_{touch}$ (top) or $R_{whisking}$ (bottom). Grey dots, other neurons. These maps exclude the ablated neurons, whose position is indicated with a faint orange background. Center, $R_{touch}$ (top) and $R_{whisking}$ (bottom) for the ablated population. Right, $R_{touch}$ (top) and $R_{whisking}$ (bottom) following ablation, with ablated neurons again excluded. **b.** As in **a**, but for silent neuron ablation.

**Extended Data 9.** Simulation of whisking neuron ablation produces representation degradation in networks with elevated, but not equal subnetwork connectivity. **a.** Whisking input was simulated by using input with a peak time of 50 ms (bottom), in contrast to 10 ms for touch (top; Fig. 1). **b.** Model network responses aligned to input prior to ablation. Left to right, increasing subnetwork connectivity. Bottom to top, raster plots (each showing a subset of neurons from an example network) and peristimulus time histograms (PSTHs; averaged across all neurons and networks) for the three neuronal populations in the model (Figure 1a; Methods). **c.** As in **b**, but following ablation. Thin PSTHs are before ablation. **d.** Effect of ablation on encoding score distributions across networks. P-values are for nested bootstrap (Methods) across multiple simulated networks. For equal subnetwork connectivity simulations ($P_{conn}$ = 0.2; top), encoding score shifted from 0.129 ± 0.011 to 0.135 ± 0.011 (grand median ± adjusted M.A.D., P = 0.003, N = 30 networks, Wilcoxon signed rank test). For elevated subnetwork connectivity simulations ($P_{conn}$ = 0.4, bottom), encoding score shifted from 0.142 ± 0.024 to 0.104 ± 0.013 (P < 0.001, N = 30 networks). Both effects are consistent with what was observed in touch stimulus simulations (Figure 1).

#### Materials and Methods

##### Network model

We modeled the L2/3 network associated with a single vS1 column, one of the most extensively studied cortical microcircuits[3], as a network of leaky integrate-and-fire neurons[4]. The dynamics of each neuron were governed by:

$$(1)\ \tau \frac{dV_i}{dt} = V_r - V(t) + R[I_i^{exc}(t) + I_i^{inh}(t) + I_i^{ext}(t)].$$

Here, $V$ is the membrane potential, $V_r$ is the rest/reset potential, $\tau$ is the membrane time constant, $R$ is the input resistance, $I^{exc}$ and $I^{inh}$ are excitatory and inhibitory synaptic currents, $I^{ext}$ is a current representing sensory stimulus drive (e.g. from layer 4 inputs), and $i$ indexes the neurons in the network. When the membrane potential reaches the spiking threshold, $V_{th}$, a spike is emitted, the membrane potential is reset to $V_r$, and the dynamics of the neuron are frozen for a short refractory period, $t_{ref}$. The synaptic currents follow kick-and-decay dynamics:

$$(2)\ \tau_{syn} \frac{dI_i^{syn}}{dt} = -I_i^{syn} + \tau_{syn} \sum_{j,k} w_{ij} \delta(t - t_{jk}^{syn} - t_d).$$

In this equation, $syn$ denotes the type of synapse (either $exc$ or $inh$), $\tau_{syn}$ is the synaptic time constant, $w_{ij}$ is a matrix of synaptic strengths from neuron $j$ to neuron $i$, $t_{jk}$ is the time of the $k^{th}$ spike of neuron $j$, and $t_d$ is the spike transmission delay. The sum over $j$ is over all neurons, while the sum over $k$ is over all spikes from that neuron.

The network comprised 2,000 neurons, of which 1,700 (85%) were excitatory and 300 (15%) were inhibitory[5]. Excitatory neurons had $\tau$ = 30 ms, whereas inhibitory neurons had faster dynamics with $\tau$ = 10 ms[5,6]. As the subthreshold dynamics for these neurons are linear, the behavior of these neurons is invariant to changes of scale in $V$. The meaningful quantity is $\Delta V = V_{th} - V_r$, which we assume to be, on average, 35 mV for all neurons[5,6]. Each neuron was assigned a value chosen uniformly within a range of ±50% of this mean value. Excitatory and inhibitory post-synaptic currents (PSCs) had time constants of $\tau_{exc}$ = 2 ms and $\tau_{inh}$ = 3 ms, respectively. Neurons had a refractory period of $t_{ref}$ = 0.5 ms. Synapses had a mean synaptic delay of $t_d$ = 0.6 ms[6]; each individual synapse was assigned a value chosen within a range of ±50% of this mean value. While all neural parameters could be modeled as random variables, we found that jittering $\Delta V$ for the neurons and $t_d$ for the synapses provided sufficient heterogeneity to give a broad range of baseline spike rates across neurons and also avoid artifactual synchronized activity. The input resistance, $R$, was factored into the synaptic weights and the magnitudes of the external currents, as described below.

The excitatory population was subdivided into two groups: a small input-recipient subnetwork of 200 neurons and the remainder of the excitatory population (1,500 neurons). Although all neurons in L2/3 likely receive touch-related input from L4[7], only a small fraction are driven to spike after touch[1,8], justifying this model assumption. Simulations where all neurons received sensory input, but subnetwork neurons received more, produced qualitatively comparable results (described below). We neglected feedforward inhibition[9]. This is justified because feedforward inhibition simply rescales input from L4[10], which in our model is adjusted to produce experimental observed population activity levels. Thus, each neuron belonged to one of three groups: the excitatory subnetwork (S), the remainder of the excitatory population (E), or the inhibitory population (I). Connections between neurons are determined by a block stochastic model. Given two neurons – the first from group A, the second from group B – there was a fixed probability of a connection from the first neuron to the second, denoted $p_{AB}$.

In our 'equal subnetwork connectivity' network, we employed sparse connectivity between excitatory neurons, $p_{SS} = p_{SE} = p_{ES} = p_{EE} = 0.2$, and more dense connectivity both within the inhibitory population as well as between the excitatory and inhibitory populations, $p_{II} = p_{IS} = p_{IE} = p_{SI} = p_{EI} = 0.6$ [5,6]. All connections had the same strength, which was chosen so that the resulting post-synaptic potential (PSP) is 1 mV[5,6]. Because it is the only connectivity parameter that varies systematically, we denote $p_{SS}$ as $P_{conn}$ in the rest of the text.

In the 'elevated subnetwork connectivity' version of the network, $P_{conn}$ was increased to 0.4. To replicate the experimentally observed relationship between connectivity and synaptic strength[11], the synaptic weight for these connections was increased to give PSPs of 1.6 mV. Networks with other levels of connectivity within the subnetwork were produced by linear interpolation/extrapolation of both $P_{conn}$ and synaptic strength between the 'equal subnetwork connectivity' and 'elevated subnetwork connectivity' cases.

The stimulus drive to the network was modeled as an external current targeting the excitatory subnetwork (group S; $I^{ext} = 0$ for all neurons in groups E and I). For each stimulus presentation, the waveform of the current was modeled with a beta distribution with shape parameters $\alpha = 3$ and $\beta = 5$. The beta distribution is defined on the interval [0, 1], giving a distinct beginning and end to the stimulus. For the chosen shape parameters, the beta distribution has a value of 0 at its endpoints and a peak at $(\alpha - 1)/(\alpha + \beta - 2)$. To model the fast touch stimulus, the waveform was stretched in time so that the peak occurs 10 ms after the start[12]. The amplitude of the stimulus waveform was chosen so that the network response matched the experimental data. All neurons also receive tonic background input in the form of a Poisson spike train of excitatory spikes with a frequency of 5,000 Hz for excitatory neurons and 2,000 Hz for inhibitory neurons; these values were selected so that the tonic firing rate of these populations were ~0.5 Hz for excitatory neurons and ~10 Hz for inhibitory neurons[13,14].

Simulations were performed in Python using the Brian2 simulation package[15] with a step-size of $dt = 0.1$ ms. For each randomly sampled network connectivity, the network was first simulated for 20 s of model time (corresponding to 66 stimulus presentations) and spike trains for all neurons were recorded. The activity of each neuron was then given an encoding score, which quantifies the signal-to-noise ratio of the representation of the stimulus: the neuron's spike train was convolved with a Gaussian kernel (standard deviation, 20 ms) to produce a firing rate; the firing rate as well as the stimulus waveform was down-sampled by a factor of 5 (to a sample period of 0.5 ms) and the normalized cross-correlation between the signals was computed for leads/lags up to 10 ms; the peak value this cross-correlation is the encoding score.

We ran an initial set of exploratory simulations to constrain the strength of the input to the subnetwork so as to match physiological data[1]. We simulated both the 'equal subnetwork connectivity' ($P_{conn} = 0.2$) and the 'elevated subnetwork connectivity' ($P_{conn} = 0.4$) networks across a range on sensory input strengths. In both cases, we examined the distributions of excitatory neuron encoding scores as a function of input strengths, selecting the input strength that produced a distribution most closely matching the experimental data (in terms of distribution shape and fraction of neurons encoding the stimulus). With input strengths defined for these two cases, we then used linear interpolation/extrapolation to select the input strength for other amounts of subnetwork connectivity. Because this procedure ensured a fixed network output following the simulated sensory stimulus, amplification (ratio of network output to sensory input; Fig. 1c) was defined as the inverse of the sensory input strength [10]. Amplification was normalized to the case where neurons within the input-recipient subnetwork had connectivity probabilities equal to the non-input recipient excitatory population (that is, $P_{conn} = 0.2$).

The final set of simulations explored the effects of targeted ablation across a range of subnetwork connectivity levels (Figs. 1,4, Extended Data 1), number of ablations (Extended Data 5), and input temporal kinetics (Extended Data 9). To simulate targeted ablation, we employed 30 randomized network instances for each subnetwork connectivity level, number of ablations, and input rise time, and calculated the pre-ablation encoding score for every neuron. The 20 excitatory neurons with the highest encoding score were then removed (all outgoing synaptic strengths set to 0), the simulation was repeated, and the post-ablation encoding scores were computed. Examination of the effects of the number of ablated neurons was done with the top 0, 10, 20, 50 and 100 neurons (Extended Data 5). This was repeated for 30 different realizations of the stochastic network connectivity. Neurons were considered to be part of the sensory input representation if they had an encoding score above 0.1; the effect of the ablation (Fig. 1e, f; Extended Data 1; Extended Data 9) was quantified as the change in encoding score across all neurons that met the 0.1 encoding score criteria either before and/or after ablations. The response similarity analysis (Fig. 4) used a more stringent encoding score cutoff of 0.25; using a cutoff of 0.1 did not alter the result, but reduced the magnitude of the observed effect. The ablated neurons were excluded for both pre-

and post-ablation encoding score distribution calculations, as well as in constructing peristimulus time histograms (Fig. 1b,e, Extended Data 1, Extended Data 9).

In modeling the effect of the number of neurons ablated on the change in encoding score (Extended Data 5), we restricted our modeling to the elevated subnetwork connectivity case, $P_{conn} = 0.4$. For simulations of 'whisking' input (Extended Data 9), we shifted from a 'touch'-like stimulus, peaking 10 ms after stimulus onset, to one peaking 50 ms after onset to mimic the response to whisking input observed *in vivo* [8,12,14,16]. We examined both the equal ($P_{conn} = 0.2$) and elevated ($P_{conn} = 0.4$) connectivity cases.

For large subnetwork connectivity ($P_{conn} > 0.4$), the network transitioned to all-or-none behavior, where strong enough input can drive the subnetwork into a state of persistent elevated firing. (Extended Data 1) [17]. To reset the network after such a transition, we introduced a strong pulse of excitatory current to the inhibitory population 300 milliseconds after the stimulus onset. In this regime the extrapolation scheme for setting the stimulus strength did not apply, since spike rate was dictated by network properties and not the input strength. Instead, the input strength affected the reliability with which a stimulus would cause a transition to the state of elevated firing. We found that choosing a stimulus strength of twice what our extrapolation scheme dictated produced a reasonable number of stimulus-encoding neurons. Thus, we used this criterion to set the input strength in this regime.

We also examined a scenario (data not shown) where the stimulus input was targeted to all excitatory neurons rather than just the subnetwork. The results were qualitatively similar in that clustering was required for the network to exhibit susceptibility to ablation. However these results were difficult to interpret, as adding even small amounts of extra subnetwork connectivity to the entire excitatory population quickly made the network unstable under all but the smallest of inputs.

Peri-stimulus time histograms (PSTHs) were constructed by averaging the Gaussian-convolved (kernel standard deviation, 20 ms) responses of individual neurons, aligned to the sensory input. Code for reproducing the simulations as well as the subsequent analyses can be found at https://github.com/jwittenbach/ablation-sim.

**Determining model synaptic weights**

Our model synapses are defined by kick-and-decay dynamics of the post-synaptic currents. We set the synaptic weights to produce a desired amplitude of the resulting unitary post-synaptic potential (PSP). Here, we derive the relationship between the synaptic weight and the PSP size that allows us to do this.

Assume that a synaptic current starts at $I = 0$ when a single spike of weight $w$ arrives at $t = 0$. In the absence of any other spikes, the subsequent time-course of the current is found by integrating (2):

(3) $I(t) = we^{-t/\tau_{syn}}$ .

To determine the resulting behavior of $V(t)$, make the assumption that the PSP evolves according to a difference of exponentials:

(4) $V(t) = V_0(e^{-t/\tau_1} - e^{-t/\tau_2})$.

Differentiating (4) and using (3), one can show that this form does indeed solve (3) if we choose:

(5) $\tau_1 = \tau_m$, $\tau_2 = \tau_{syn}$, and $V_0 = \dfrac{Rw}{\tau_m / \tau_s - 1}$.

The PSP size is the maximum value of $V(t)$, which we can compute as:

(6) $V_{\max} = V_0 a^{(1-a^{-1})^{-1}}(a-1)$,

where $a = \tau_m / \tau_s$ is the ratio of the time constants. Comparing this to the expression for $V_0$ in (6) we find our desired relationship:

(7) $Rw = V_{\max} a^{(1-a^{-1})^{-1}}$.

The linearity of the synaptic dynamics allows us to use (7) to avoid explicitly determining the value of $R$.

**Animals**

All procedures were performed in compliance with the Janelia Research Campus Institutional Animal Care and Use Committee and the New York University University Animal Welfare Committee. Two transgenic lines were used for these experiments, differentiated in Extended Data 1 by a 'j' or 'n' in Animal ID. Animals whose ID starts with the letter 'j' consisted of mice expressing nuclear localized mCherry in a Cre-dependent manner[1] (R26-LSL-H2B-mCherry; JAX 023139) crossed with mice expressing Cre in cortical pyramidal neurons,[18] (Emx1-IRES-Cre; JAX 005628). In cortical L2/3, these animals expressed nuclear mCherry only in nuclei of excitatory neurons. Multiple (9 or 12) injections (450 μm deep, 300 μm apart; beveled pipettes, World Precision Instruments; 20 nL each, at 10 nL/min with oil microinjector, Narishige) of AAV2/1-syn-GCaMP6s (UPenn AV-1-PV2824) [19] were made in vS1 (3.6 mm lateral, 1.5 mm posterior) of young adult mice[1] (6-8 week). Following viral injection, a titanium headbar was attached to the skull and the craniotomy was covered with a cranial window. Animals whose ID starts with the letter 'n' consisted of mice expressing GCaMP6s in a Cre-dependent manner (Ai162[2], JAX 031562), crossed with Slc17a7-IRES2-Cre[2] (JAX 023527) to restrict expression to in pyramidal neurons. Surgeries for these animals did not include viral injections but were otherwise identical.

**Behavior**

Approximately one week after surgery, mice were trimmed to a single row of whiskers (typically the C row) and placed on water restriction[20] (1 mL per day). Training commenced 5-7 days after restriction onset. Animals were trained on an object localization task (Fig. 2) [21,22]. If the pole appeared in a range of proximal positions, the animal would be rewarded with a small water drop for licking the right ('contralateral') lickport; pole presentation at the distal position would be rewarded upon licking the left ('ipsilateral') lickport (Fig. 2a). Trials consisted of a 1-1.2 s sample epoch followed by a 0.5-1.2 s delay epoch after which a 50 or 100 ms 3.4 kHz auditory response cue signaled to the animal to respond (Fig. 2b). To prevent premature licking, the lickport was brought into tongue range by a motor (Zaber) only during the response epoch (Fig. 2b). Mice exhibiting excessive premature licking (licks before reward cue on >20% of trials; licks monitored with a laser beam; Thorlabs) were not used.

Mice were trimmed to a single whisker after reaching criterion performance (d' > 1.5 for two consecutive days). The spared whisker barrel column was identified using the neuropil signal, as described previously[1. Whisker videography was performed at 400-500 Hz. Whiskers were tracked using an automated software pipeline{Clack, 2012 #7290] and then curated using custom browser software[1].

**Imaging**

Calcium imaging was performed using a custom two-photon microscope (http://openwiki.janelia.org/wiki/display/shareddesigns/MIMMS) with a 16X, 0.8 NA objective (Nikon) objective[1]. GCaMP (BG22; Chroma) and mCherry (675/70 filter; Chroma) fluorescence was imaged using GaAsP PMTs (Hamamatsu). The 940 or 1000 nm (Coherent) imaging beam was steered with a 16 kHz line rate resonant galvanometer (Thorlabs); a piezo collar (Physik Instrumente) moved the focus axially. 512 x 512 pixel images were collected at 7 Hz, with three 600 x 600 µm images per piezo cycle. Planes were spaced either 15 µm apart. Scanimage[23] (Vidrio Technologies, http://www.vidriotech.com) controlled the microscope. Three planes, constituting a 'subvolume', were imaged simultaneously (4-6 subvolumes, spanning 45 µm each, 12-18 total planes, 180-270 µm in depth total), and power was modulated with depth using a length constant of 250 µm. Deeper subvolumes were typically imaged with higher power, using a similar length constant. Individual subvolumes were typically imaged for 50 trials per day, with all subvolumes usually visited on any given day. Alignment across days was performed as described previously[1,24].

Imaging data were processed using a semi-automated software pipeline that included image registration, segmentation, neuropil subtraction, $\Delta F/F$ computation, and event detection[1]. The de-noised $\Delta F/F$ trace, which consisted of the event amplitude trace convolved with event-specific exponential rise and decay, was used for analysis.

**Neuronal classification**

Neurons were classified using a linear-nonlinear encoding model[1,25]. The model consisted of a cascaded generalized linear model that used a temporal and stimulus domain kernel to predict the activity of individual neurons given whisker angle, θ, or whisker curvature, κ, assuming Gaussian noise with input nonlinearities.

The model predicted the neuronal response, *r* (i.e., ΔF/F), as

$$r \sim Norm(z, \sigma^2)$$
$$z = f_1(s_1) * k_1 + f_2(s_2) * k_2$$

Here, $s_1$ and $s_2$ are the whisker angle, θ, and whisker curvature, κ, respectively. The terms $f_1$ and $f_2$ are static, point-wise nonlinearities comprising a weighted sum of sixteen triangular basis functions

$$f = \sum_{i=1}^{16} w_i b_i(x)$$

Where x is the input ($s_1$ or $s_2$), with each $b_i$ given by

$$b_i = \begin{cases} (x - x_{i-1})/(x_i - x_{i-1}), & i > 1,\ x_{i-1} < x < x_i \\ (x_{i+1} - x)/(x_{i+1} - x_i), & i < N,\ x_i \le x < x_{i+1} \\ 0, & \text{otherwise} \end{cases}$$

$k_1$ and $k_2$ are temporal kernels consisting of 14 time points (2 seconds).

Thus, the model gives a z-scored prediction of neural activity, *z*, by fitting parameters $k_1$, $k_2$, $f_1$, $f_2$ for given whisker kinematic parameters, $s_1$ and $s_2$.

The model parameters $k_1$, $k_2$, $f_1$, $f_2$ were fit using maximum likelihood with block coordinate descent. This procedure reliably estimated model parameters within three to five iterations.

To avoid degeneracy associated with arbitrary scaling factors on either the kernels or the nonlinearities, the nonlinearities were forced to have minimum of 0 and maximum of 1. Temporal kernels were unconstrained.

Because whisker movements were sampled at 500 Hz while calcium responses were measured at 7 Hz, the nonlinearity was applied to the whisker variables at their native temporal resolution, followed by linear down sampling to 7 Hz. Thus, additional information in the higher resolution whisker variables was incorporated into the prediction.

A prior was used to ensure smoothness of both the temporal kernels and the nonlinearities and prevent over-fitting. Specifically, the prior added a factor to the objective function penalizing excessive second derivatives of the temporal kernels and nonlinearities. Employing such a prior corresponds to maximizing the log-posterior, with the prior adding a small penalty to the objective function. In order to fit several thousand cells efficiently, the scale factor associated with this penalty was determined from a cross-validated inspection of a random subset of neurons.

The model was fit using five-fold cross-validation across trials. That is, a randomly selected 80% of trials were used for fitting, and 20% were used for evaluation, with 5 distinct evaluation groups per fit ensuring all data was used for fitting in exactly one case. The Pearson correlation between the predicted and actual $\Delta$F/F yielded a measure of the quality of the model fit. This correlation was used as the encoding score for vS1 data: $R_{touch}$, based on $\Delta\kappa$ for touch, and $R_{whisking}$, based on whisker $\theta$ for whisking (Fig. 2d,e). A neuron was considered part of a representation if $R_{touch}$ or $R_{whisking}$ exceeded 0.1 and if the neuron's score was above the 99$^{th}$ percentile of scores measured from matched shuffled neural activity. These criteria are more stringent than those employed previously[1], because ablation predominantly impacted neurons with high encoding scores. Using a less stringent criterion did not change the underlying conclusions, but did dilute the magnitude of the effect.

Shuffled activity was generated by randomizing the timing of the calcium events to construct a novel de-noised $\Delta$F/F trace. Matched shuffled activity was selected by using neurons from the same imaging subvolume (i.e., concurrently imaged to ensure identical animal behavior) sharing a similar event rate. Event rates were matched by partitioning the neurons from a subvolume into 10 equally sized bins (in terms of neuron count). Thus, a given neuron's R had to exceed the 99$^{th}$ percentile of R's obtained across neurons in the same subvolume and event rate bin whose event times were shuffled.

We examined robustness by partitioning individual days into two interdigitated pseudo-sessions, and measuring the correlation between encoding scores for the two pseudo-sessions[1]. The resulting correlations ranged between ~0.5-0.75. Neural classification (whisking and touch) was stable across days for trained mice. Furthermore, the touch neuron curvature kernels assumed 'V' or 'L'-like shapes[1], meaning that high magnitude curvature changes drove the largest responses, a result consistent with the known responses of these neurons. Finally, the kernels were stable across days[1]. Thus, changes in vS1 encoding following ablation are not a reflection of the variability inherent to the encoding model, but rather reflect genuine changes in the underlying representations.

Ablated neurons were always excluded from analysis of experimental data, including calculations of pre-ablation population encoding scores. Analyses involving 'pre-ablation' and 'post-ablation' data were pooled across multiple behavioral sessions. Pre- and post-ablation data each consisted of at least 2 (but typically 3) behavioral sessions, with each subvolume sampled on most sessions. Given that a single subvolume was imaged for around 50 trials in a session, classification typically employed around 150 behavioral trials per neuron (minimum for encoding model: 100 trials). Neurons participating in both representations in a given area were excluded from analysis. In all cases where comparisons of pre- and post-ablation distributions were made, neurons were included for analysis if they met the criteria for representation membership (described above) in the pre- or post-ablation period, or during both periods.

**Multiphoton ablation**

Ablations [26-28] were performed with 880 nm (Chameleon Ultra 2; Coherent) or 1040 nm (Fidelity HP; Coherent) femtosecond laser pulses delivered through a 0.8 NA, 16 X objective (Nikon) in animals that were awake but not performing the task or lightly anesthetized using isoflurane (1-2%). Cells were selected for ablation using brain region specific criteria applied to imaging data from the first three behavioral sessions. On the day of ablation, a new image was acquired and a warp field transform of was employed to find the selected neurons[24]. Imaging was not performed on the day of ablation; post-ablation analyses used the imaging data from the 2-4 behavioral sessions on the days following ablation. Typically, experiments consisted of three days of pre-ablation data collection, the ablation day, and three days of post-ablation data collection.

In virally transfected mice, the mCherry signal was used to restrict ablations to pyramidal neurons[1]; in mice endogenously expressing GCaMP, presence of fluorophore was used for this purpose. The neurons with strongest touch or whisking encoding scores within the spared barrel column were targeted for ablation. The percentage of touch and whisking neurons ablated was calculated in relation to the estimated 1,691 pyramidal neurons in L2/3 [5], of which 17% (287) belonged to each representation[1]. Neurons participating in both representations (touch and whisking) were avoided. For vS1 silent cell ablation, neurons with a calcium event rate below 0.025 Hz were targeted for ablation.

In all cases, approximately 2/3 of ablations were successful (Extended Data 3c). Proximity to vasculature, low baseline fluorescence, and excessive depth accounted for most failures. Thus, despite targeting the strongest neurons in a given representation, the actual representation strength of ablated neurons varied. The encoding score percentile of the ablated neurons is therefore reported in the text.

After ablation we consistently observed an increase in GCaMP6 fluorescence in the targeted neuron (Extended Data 3a,b). Taking advantage of this signature, the ablation protocol consisted of interleaved 'ablation' and 'evaluation' epochs (Extended Data 2b; ablation duration: 50-200 ms; evaluation: 0.1-2 s, with longer evaluation times proving more reliable). Ablation epoch power started at the evaluation epoch power (25-100 mW, measured at the specimen) and rose linearly as necessary (up to 1W) over the course of several seconds. The beam was focused on the brightest part of the targeted neuron using a pair of galvanometers (Cambridge Technology) and oscillated over a path spanning 1-2 μm. Evaluation epoch fluorescence data was collected using standard resonant galvanometer imaging[1], though restricted to a single plane (~30 Hz). Ablations were terminated after observation of the fluorescence rise in the target neuron.

Our protocol could effectively remove 2/3 of targeted neurons (Extended Data 3c-e). Consistent with similar protocols[29,30], ablation did not produce off-target damage:

calcium event rates for neurons adjacent (10-25 µm) to ablated cells did not change following the ablation of silent neurons (pre-ablation event rate: 0.014 ± 0.008 Hz, grand median ± adjusted M.A.D., Methods, post-ablation: 0.014 ± 0.007 Hz; P = 0.721 pre- vs. post-ablation, Wilcoxon signed rank test, N = 8 mice; 1,028 neurons across all mice), and glial immunoreactivity was confined to the site of the lesioned neuron (Extended Data 4).

Typically, 10-50 ablations were performed over the course of one hour. For histological analysis (Extended Data 4b-d), perfusion was performed 24h after ablation in two Emx1-Cre X LSL-H2B-mCherry animals (9 and 28 ablations were successful in these animals, approximating typical experimental conditions). Alternating cryomicrotome (Leica) sections were exposed to antibodies for either the microglial marker iba1 (Abcam, ab5076) or the astrocytic marker GFAP (Abcam, ab7260). Glial reactions were measured by first locating the ablated neuron's center in the glial immunoreaction image. An edge detection algorithm[19] that operated on an intensity image in angle-distance space from the neuron center detected the reaction's extent (Extended Data 4d). Specifically, intensity profiles were measured across a range of angles emanating from the point within the ablation. To delimit the glial reaction, large drops in intensity were detected. Glia was considered reactive if the intensity inside the detected reaction area was two standard deviations above background image intensity.

Following ablation, nearby neurons retained sensory responses (Fig. 2f), event rate (Extended Data 4a), and maintained structural integrity (Extended Data 4b,c,e). In a few instances (N = 5) the ablation termination protocol failed, resulting in more extensive lesions (Extended Data 3f). These experiments were excluded from the study.

We excluded ablated neurons as well as neurons within a cylinder centered on ablated neurons having a radius of 10 µm and a height of 60 µm from all analyses. This excluded neurons abutting ablated cells, and ensured no neurons within a typical glial reaction radius would be included.

**Response similarity**

For simulated data, response similarity was measured by taking the Pearson correlation of the Gaussian-convolved (kernel standard deviation, 20 ms) activity of an individual neuron with the mean Gaussian-convolved activity of the ablated neurons (Fig. 4a). For experimental data, response similarity was measured by correlating the individual neuronal trial averaged ΔF/F to the ablated neuron trial average mean. For each neuron, trial averaged ΔF/F was calculated by taking the mean ΔF/F across all correct lick contra and ipsi trials (Fig. 4c), then concatenating these two vectors (Fig. 4d). The mean of these vectors across the ablated neurons constituted the 'ablated neuron mean' (Fig. 4d). Response similarity is simply the Pearson correlation of the individual neuron trial averaged ΔF/F with the mean trial averaged ΔF/F across all ablated neurons. Single-network (Fig. 4b) or animal (Figs. 4e, f, g) averages were computed with response similarity bins having a width of 0.1. The grand mean of these is shown as a dark line on these plots.

Trial-averaged ΔF/F correlations were employed instead of raw activity correlations because neurons were not all imaged simultaneously; only neurons in a given subvolume were imaged simultaneously. Because ablated neurons came from multiple subvolumes, response similarity had to employ trial-averaged responses to allow for comparison across disjointly recorded populations.

**Statistical analyses**

The majority of statistical comparisons were performed using the Wilcoxon signed rank test comparing paired medians within individual animals for two conditions (e.g., before and after ablation, or proximal and distal encoding score change). For cases where values had no natural pairing (comparison of different ablation types), the Wilcoxon rank sum test comparing medians was used to compare distributions. To test if encoding score change depended on response similarity (Fig. 4), we first fit a line to individual networks or animals (e.g., Fig. 4e; the linear fit is distinct from the cross-cell mean that is shown). Next, we tested whether the slopes thus obtained were distinct from 0 across all networks or animals using the non-parametric sign test. In all cases, we used the median of single-neuron values within an animal. That is, we treated mice, and never neurons, as independent observations. Where given, adjusted median absolute deviation (M.A.D.) was calculated by multiplying the median absolute deviation by 1.4826 so as to approximate the standard deviation under conditions of normality.

**References:**


1 Peron, S. P., Freeman, J., Iyer, V., Guo, C. & Svoboda, K. A Cellular Resolution Map of Barrel Cortex Activity during Tactile Behavior. *Neuron* **86**, 783-799, doi:10.1016/j.neuron.2015.03.027 (2015).

2 Daigle, T. L. *et al.* A Suite of Transgenic Driver and Reporter Mouse Lines with Enhanced Brain-Cell-Type Targeting and Functionality. *Cell* **174**, 465-480 e422, doi:10.1016/j.cell.2018.06.035 (2018).

3 Petersen, C. C. & Crochet, S. Synaptic computation and sensory processing in neocortical layer 2/3. *Neuron* **78**, 28-48, doi:10.1016/j.neuron.2013.03.020 (2013).

4 Brette, R. *et al.* Simulation of networks of spiking neurons: a review of tools and strategies. *J Comput Neurosci* **23**, 349-398, doi:10.1007/s10827-007-0038-6 (2007).

5 Lefort, S., Tomm, C., Floyd Sarria, J. C. & Petersen, C. C. The excitatory neuronal network of the C2 barrel column in mouse primary somatosensory cortex. *Neuron* **61**, 301-316, doi:10.1016/j.neuron.2008.12.020 (2009).

6 Avermann, M., Tomm, C., Mateo, C., Gerstner, W. & Petersen, C. C. Microcircuits of excitatory and inhibitory neurons in layer 2/3 of mouse barrel cortex. *J Neurophysiol* **107**, 3116-3134, doi:10.1152/jn.00917.2011 (2012).

7 Bureau, I., von Saint Paul, F. & Svoboda, K. Interdigitated Paralemniscal and Lemniscal Pathways in the Mouse Barrel Cortex. *PLoS Biol* **4**, e382 (2006).

8 Crochet, S., Poulet, J. F., Kremer, Y. & Petersen, C. C. Synaptic mechanisms underlying sparse coding of active touch. *Neuron* **69**, 1160-1175, doi:10.1016/j.neuron.2011.02.022 (2011).

9 Xue, M., Atallah, B. V. & Scanziani, M. Equalizing excitation-inhibition ratios across visual cortical neurons. *Nature* **511**, 596-600, doi:10.1038/nature13321 (2014).

10 Chance, F. S., Nelson, S. B. & Abbott, L. F. Complex cells as cortically amplified simple cells. *Nat Neurosci* **2**, 277-282, doi:10.1038/6381 (1999).

11 Biagi, R., Cossellu, G., Sarcina, M., Pizzamiglio, I. T. & Farronato, G. Laser-assisted treatment of dentinal hypersensitivity: a literature review. *Annali di stomatologia* **6**, 75-80, doi:10.11138/ads/2015.6.3.075 (2015).

12 Hires, S. A., Gutnisky, D. A., Yu, J., O'Connor, D. H. & Svoboda, K. Low-noise encoding of active touch by layer 4 in the somatosensory cortex. *Elife* **4**, doi:10.7554/eLife.06619 (2015).

13 Gentet, L. J. *et al.* Unique functional properties of somatostatin-expressing GABAergic neurons in mouse barrel cortex. *Nat Neurosci* **15**, 607-612, doi:10.1038/nn.3051 (2012).

14 Yu, J., Gutnisky, D. A., Hires, S. A. & Svoboda, K. Layer 4 fast-spiking interneurons filter thalamocortical signals during active somatosensation. *Nat Neurosci* **19**, 1647-1657, doi:10.1038/nn.4412 (2016).

15 Stimberg, M., Brette, R. & Goodman, D. F. Brian 2, an intuitive and efficient neural simulator. *Elife* **8**, doi:10.7554/eLife.47314 (2019).

16 Curtis, J. C. & Kleinfeld, D. Phase-to-rate transformations encode touch in cortical neurons of a scanning sensorimotor system. *Nat Neurosci* **12**, 492-501 (2009).

17 Litwin-Kumar, A. & Doiron, B. Slow dynamics and high variability in balanced cortical networks with clustered connections. *Nat Neurosci* **15**, 1498-1505, doi:10.1038/nn.3220 (2012).

18 Gorski, J. A. *et al.* Cortical excitatory neurons and glia, but not GABAergic neurons, are produced in the Emx1-expressing lineage. *The Journal of neuroscience : the official journal of the Society for Neuroscience* **22**, 6309-6314, doi:20026564 (2002).

19 Chen, T. W. *et al.* Ultrasensitive fluorescent proteins for imaging neuronal activity. *Nature* **499**, 295-300, doi:10.1038/nature12354 (2013).

20 Guo, Z. V. *et al.* Procedures for behavioral experiments in head-fixed mice. *PLoS One* **9**, e88678, doi:10.1371/journal.pone.0088678 (2014).

21 O'Connor, D. H. *et al.* Vibrissa-based object localization in head-fixed mice. *J Neurosci* **30**, 1947-1967, doi:10.1523/JNEUROSCI.3762-09.2010 (2010).

22 Guo, Z. V. *et al.* Flow of cortical activity underlying a tactile decision in mice. *Neuron* **81**, 179-194, doi:10.1016/j.neuron.2013.10.020 (2014).

23 Pologruto, T. A., Sabatini, B. L. & Svoboda, K. ScanImage: Flexible software for operating laser-scanning microscopes. *BioMedical Engineering OnLine* **2**, 13 (2003).

24 Huber, D. *et al.* Multiple dynamic representations in the motor cortex during sensorimotor learning. *Nature* **484**, 473-478, doi:10.1038/nature11039 (2012).

25 Ahrens, M. B., Paninski, L. & Sahani, M. Inferring input nonlinearities in neural encoding models. *Network* **19**, 35-67, doi:10.1080/09548980701813936 (2008).

26 Konig, K., Becker, T. W., Fisher, P., Riemann, I. & Halbhuber, K. J. Pulse-length dependence of cellular response to intense near-infrared laser pulses in multiphoton microscopes. *Optics Lett.* **24**, 113-115 (1999).

27 Orger, M. B., Kampff, A. R., Severi, K. E., Bollmann, J. H. & Engert, F. Control of visually guided behavior by distinct populations of spinal projection neurons. *Nat Neurosci* **11**, 327-333, doi:10.1038/nn2048 (2008).

28 Allegra Mascaro, A. L., Sacconi, L. & Pavone, F. S. Multi-photon nanosurgery in live brain. *Frontiers in neuroenergetics* **2**, doi:10.3389/fnene.2010.00021 (2010).

29 Canty, A. J. *et al.* In-vivo single neuron axotomy triggers axon regeneration to restore synaptic density in specific cortical circuits. *Nat Commun* **4**, 2038, doi:10.1038/ncomms3038 (2013).

30 Vladimirov, N. *et al.* Brain-wide circuit interrogation at the cellular level guided by online analysis of neuronal function. *Nat Methods* **15**, 1117-1125, doi:10.1038/s41592-018-0221-x (2018).

### Peron et al., Extended Data 1

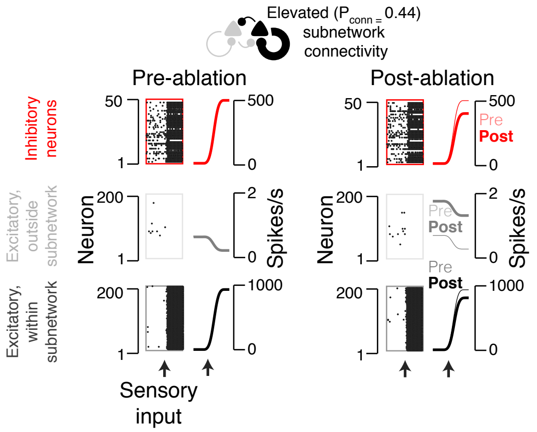

Peron et al., Extended Data 3

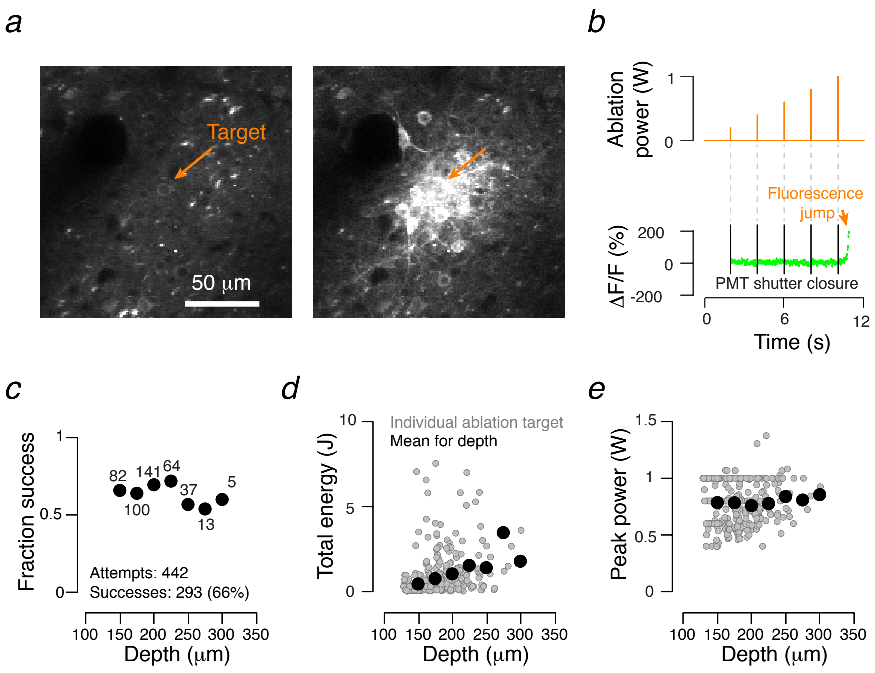

### Peron et al., Extended Data 4

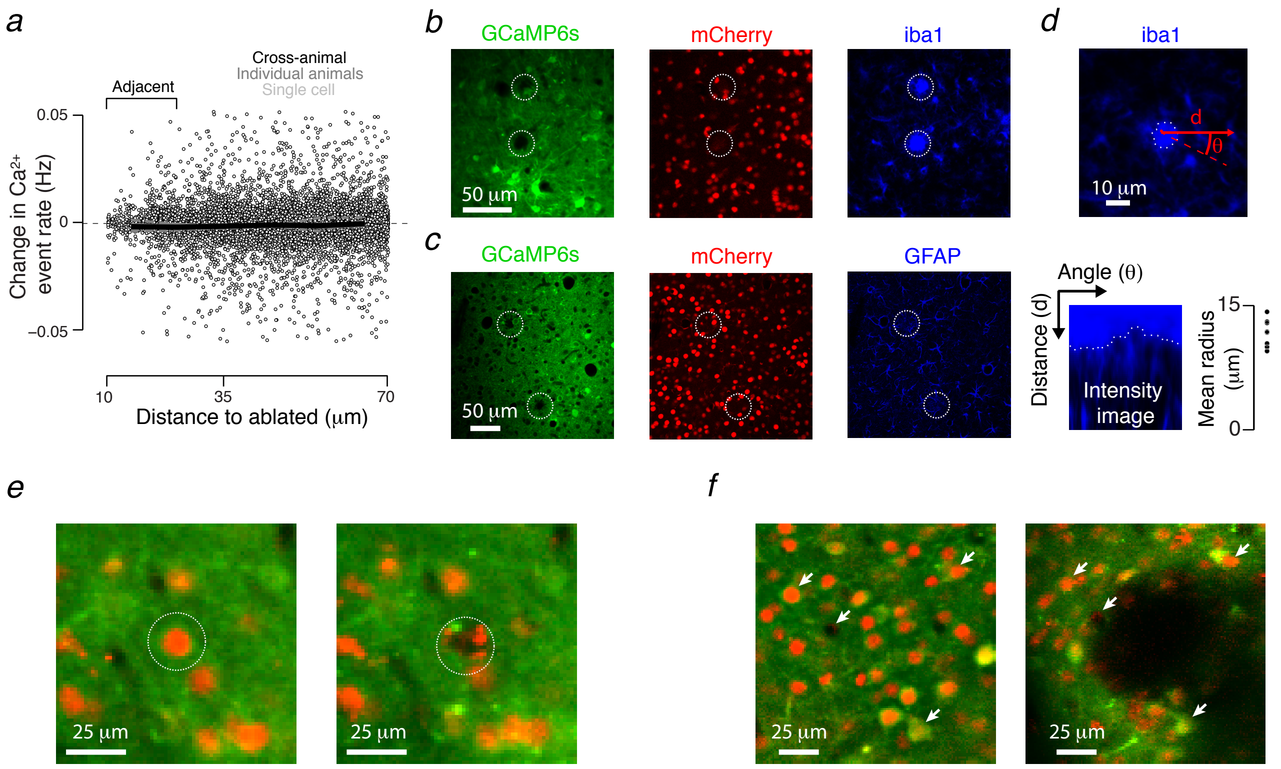

Peron et al., Extended Data 5

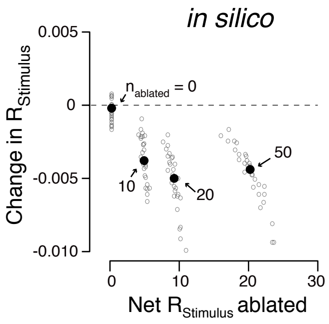

### Peron et al., Extended Data 6

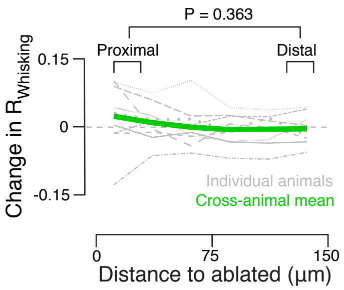

Peron et al., Extended Data 7

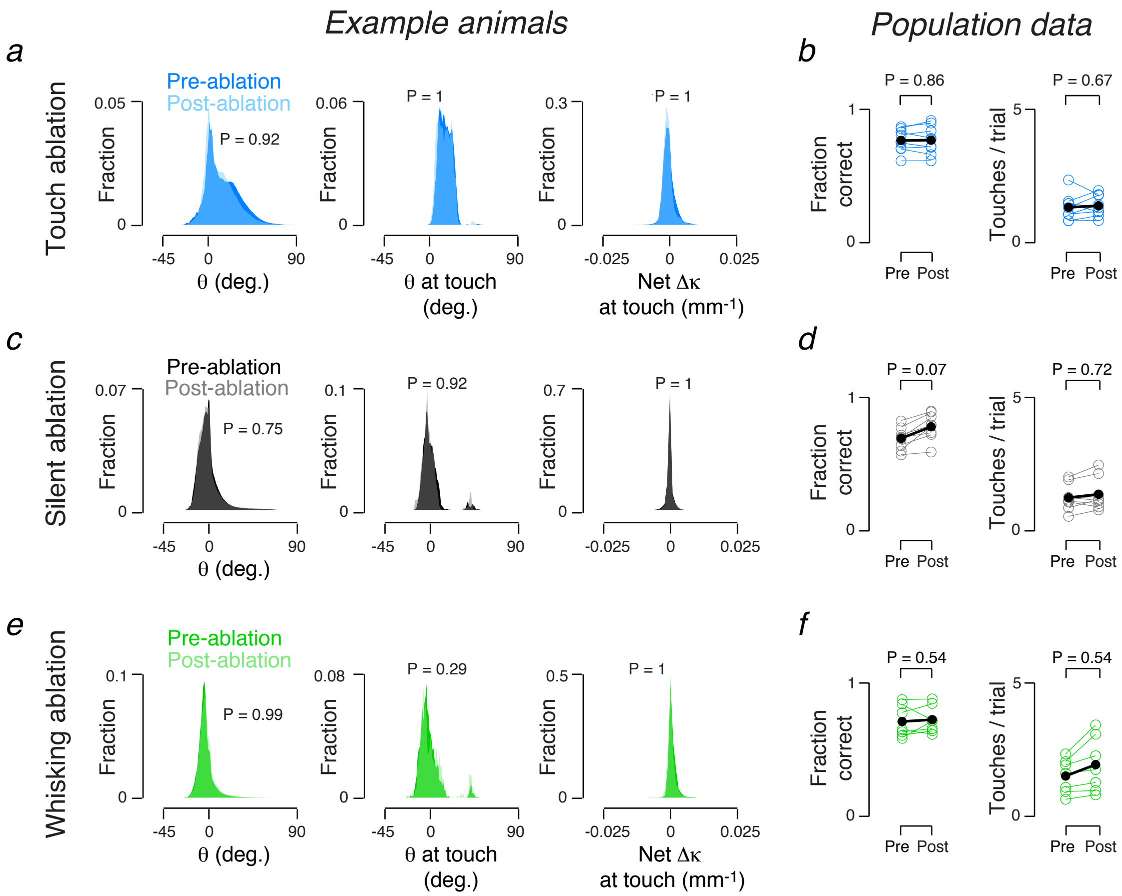

Peron et al., Extended Data 8

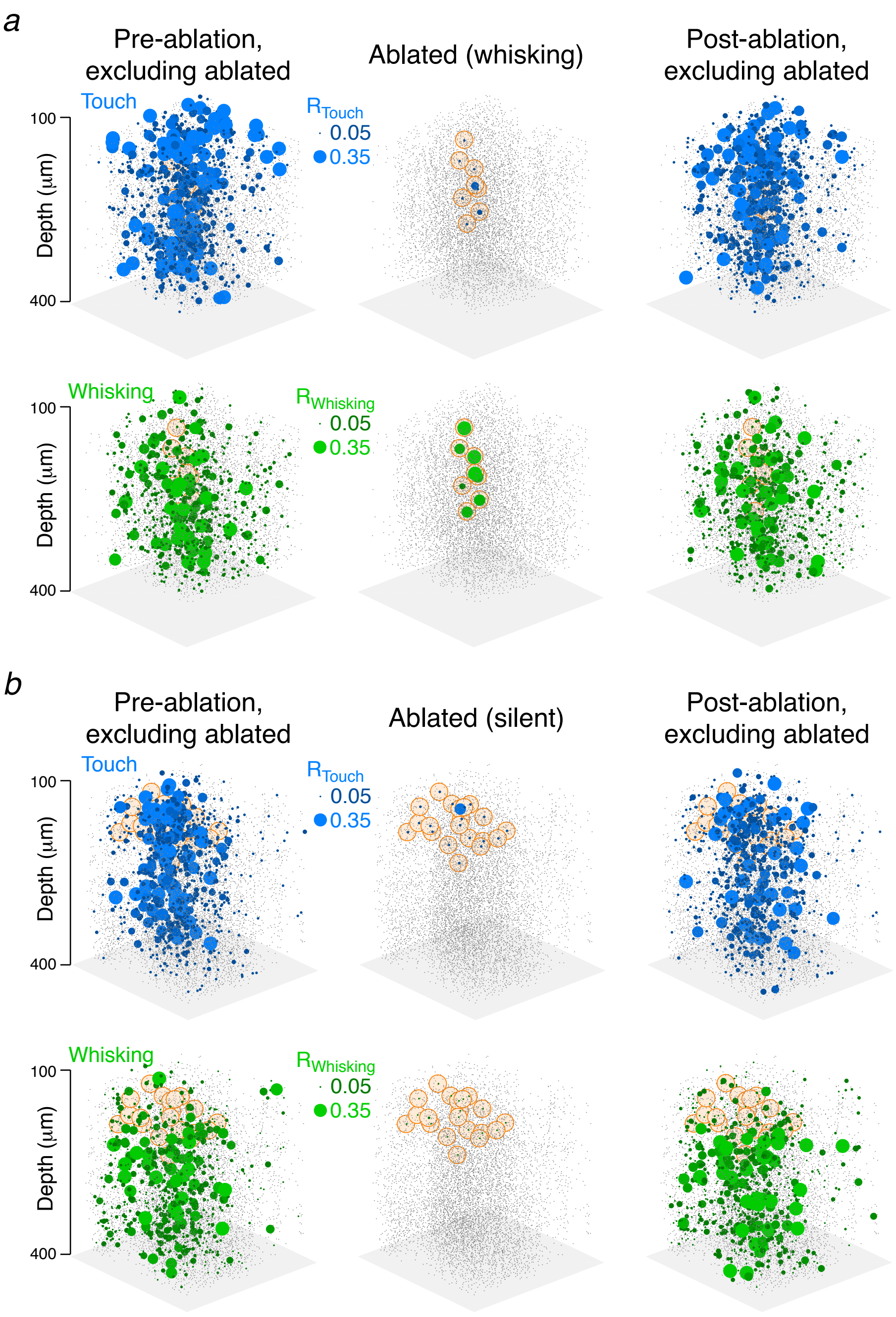

Peron et al., Extended Data 9

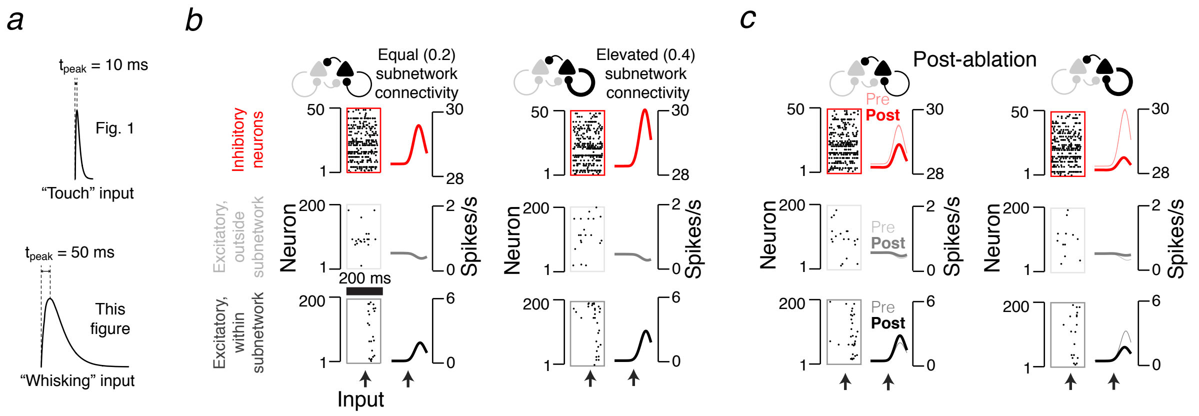
